# Representation of event and object concepts in ventral anterior temporal lobe and angular gyrus

**DOI:** 10.1101/2023.10.13.562253

**Authors:** Yueyang Zhang, Wei Wu, Daniel Mirman, Paul Hoffman

**Author notes:** Correspondence to: Dr. Paul Hoffman, School of Philosophy, Psychology & Language Sciences, University of Edinburgh, 7 George Square, Edinburgh, EH8 9JZ, UK.

## Abstract

Semantic knowledge includes understanding of objects and their features and also understanding of the characteristics of events. The hub-and-spoke theory holds that these conceptual representations rely on multiple information sources that are integrated in a central hub in the ventral anterior temporal lobes (vATL). Dual-hub theory expands this framework with the claim that the vATL hub is specialized for object representation, while a second hub in angular gyrus (AG) is specialized for event representation. To test these ideas, we used RSA, univariate and PPI analyses of fMRI data collected while participants processed object and event concepts (e.g., ‘an apple’, ‘a wedding’) presented as images and written words. RSA showed that AG encoded event concept similarity more than object similarity, although the left AG also encoded object similarity. Bilateral vATLs encoded both object and event concept structure, and left vATL exhibited stronger coding for events. PPI analysis revealed greater connectivity between left vATL and right pMTG, and between right AG and bilateral ITG and middle occipital gyrus, for event concepts compared to object concepts. These findings support the specialization of AG for event semantics, though with some involvement in object coding, but do not support vATL specialization for object concepts.

## Introduction

Humans can recognise and reason about single objects, and we can also understand events as coherent conceptual units --complex, context-bound interactions between objects that unfold over time. Object similarity can be captured by shared features, whereas events involve multiple objects’ interactions, temporal sequences and causal relationships (Altmann & Ekves, 2019). A core function of the semantic system is to represent similarities between these abstract conceptual units. For example, apples are more similar to tomatoes than to hammers, and weddings are more similar to parties than to fights. The neural coding of object similarity has been studied in depth (Bi et al., 2016; Chen et al., 2016; Devereux et al., 2013; Hutchison et al., 2014; Kaneshiro et al., 2015; Xu et al., 2018). Event structure coding is also investigated by some studies (Baldassano et al., 2017; Bedny et al., 2014; Morton et al., 2020). However, object representation and event representation are rarely compared directly, meaning that differences in their neural bases remain unclear. Thus, in the present study, we used representational similarity analysis (RSA), univariate fMRI analyses, and functional connectivity analyses to directly compare how the semantic structures of objects and events are represented in the brain.

Vision is crucial for identifying objects and events, thus specializations for object and event understanding could be driven by the organization of the visual system into dorsal and ventral pathways (Mirman et al., 2017). The dorsal pathway usually refers to the processing stream that lies between early visual cortex and frontal-parietal regions specialized for action, and which courses through temporal-parietal cortex (Kravitz et al., 2013; Mishkin et al., 1983). The dorsal stream is identified as a ‘where/how’ pathway, supporting visually-guided action, motion and spatial cognition (Andersen & Cui, 2009; Buxbaum & Kalénine, 2010; Husain & Nachev, 2007; Wager & Smith, 2003; Watson & Chatterjee, 2011). The dorsal stream may be particularly important for event representation, as this requires processing of objects’ interactions and their spatiotemporal relations. Conversely, the ventral pathway lies between early visual cortex and the ventral anterior temporal lobe (vATL), and courses through the inferior parts of the temporal lobe (Kravitz et al., 2013; Mishkin et al., 1983). This stream is characterised as a ‘what’ pathway, specialised for identifying and categorizing objects. In line with this view, ventral pathway regions are engaged in processing and integrating perceptual features like colour, size, and brightness (Baron et al., 2010; Coutanche & Thompson-Schill, 2015; Martin et al., 2018). Regions in this pathway show category-selective effects for different object categories like tools, animals and human faces (Bi et al., 2016; Hutchison et al., 2014).

As the junction of the ventral pathway with other processing streams, vATL is thought to act as a transmodal semantic hub that combines visual features with multimodal information sources to generate conceptual representations (for review, see Lambon Ralph et al., 2017). The ATLs are strongly associated with integrating object features across sensory modalities (Coutanche & Thompson-Schill, 2015; Rogers & McClelland, 2004), and are engaged in semantic processing irrespective of input modality (e.g., words, pictures and sounds) (Binney et al., 2010; Marinkovic et al., 2003; Vandenberghe et al., 1996; Visser & Lambon Ralph, 2011) and across a range of conceptual categories (Conca et al., 2021; Hoffman et al., 2015; Rice et al., 2018; Wang et al., 2019).

Studies using multivariate pattern analysis indicate that ATL regions code semantic relationships between objects (Chen et al., 2016; Fairhall & Caramazza, 2013; Peelen & Caramazza, 2012; Rogers et al., 2021). For example, in an iEEG study using a picture-naming task, Chen et al. (2016) observed that vATL activity patterns were predicted by semantic similarity between objects, even after controlling for visual and phonological features of the stimuli. The medial part of vATL, the perirhinal cortex, has been implicated specifically in recognizing objects and in differentiating between objects that have many overlapping semantic features (for review, see Clarke & Tyler, 2015). Perirhinal cortex activation increases when participants recognize semantically more-confusable objects (Clarke & Tyler, 2014; Tyler et al., 2013) and damage to this region results in deficits for naming semantically more-confusable objects (Wright et al., 2015). RSA analyses of fMRI data indicate that more similar objects elicit more similar patterns of activation in the perirhinal cortex (Bruffaerts et al., 2013; Devereux et al., 2018; Liuzzi et al., 2015; Naspi et al., 2021). For example, Liuzzi et al. (2015) presented people with written object names, and found that in left perirhinal cortex, activation pattern similarity was predicted by semantic similarity between objects (measured in terms of their property overlap). However, while it is now well-established that regions within vATL code semantic similarity between objects, it remains unclear whether this region also codes semantic similarities between events. Studies of event semantics have instead focused on regions within the temporoparietal cortex (TPC).

An association between TPC and event representation has been suggested by many researchers (for review, see Binder & Desai, 2011; Mirman et al., 2017). Event representations require frequent processing of interactions or contextual associations (e.g., action, spatial, temporal information). This kind of processing may be well-suited to TPC regions, which participate in, and receive inputs from, the dorsal visual stream. TPC regions have been implicated in the semantics of action and in representing thematic relationships between concepts. Posterior temporal lobe is involved in understanding action concepts (Bedny et al., 2014; Kable et al., 2005; Kable et al., 2002) and motion concepts (Bedny et al., 2008; Gennari et al., 2007; Noppeney et al., 2005; Watson et al., 2013; Zhang et al., 2022). The posterior parietal cortex is involved in action planning (for reviews, see Andersen & Cui, 2009; Buxbaum & Kalénine, 2010). Parietal regions within TPC are also important for integrating spatially distributed objects into a single coherent percept (Huberle & Karnath, 2012; Lestou et al., 2014) and for making temporal order judgements (Davis et al., 2009).

These roles in supporting the dynamic aspects of semantics make TPC particularly suited to representing interactions between objects. Indeed, based on neuropsychological and neuroimaging evidence, the dual-hub theory of semantic representation proposes that TPC is specialised for coding thematic/event-based semantic relations (e.g., dog-bone) and the ATL for taxonomic/similarity-based semantic relations (e.g., dog-cat) (Mirman et al., 2017; Schwartz et al., 2011). A recent fMRI meta-analysis provided support for this idea by revealing that TPC regions are reliably more activated by thematic than taxonomic relations (Zhang et al., 2023).

Within TPC, the angular gyrus (AG) in particular has been identified as a critical area for multiple functions relevant to event representation: autobiographical memory and episodic memory (Bonnici et al., 2018; Russell et al., 2019), retrieval of multimodal spatiotemporal memories (Ben-Zvi et al., 2015; Bonnici et al., 2016; Richter et al., 2016; Yazar et al., 2014, 2017), and combinatorial semantics (e.g., computing the meanings of noun+noun and verb+noun phrases) (Boylan et al., 2015; Price et al., 2015). More broadly, AG is a key part of the default mode network (DMN), which is implicated in coding situation models of ongoing events and segmenting experiences into separate events (Baldassano et al., 2017; Morales et al., 2022; Ranganath & Ritchey, 2012; Swallow et al., 2011; Yeshurun et al., 2021; Zacks et al., 2010). DMN appears to act as a dynamic network that combines incoming external information with internal information from prior experiences to create detailed, context-specific representations of situations as they develop over time (for review, see Ranganath & Ritchey, 2012; Yeshurun et al., 2021). In line with these functions, DMN is sensitive to event boundaries in a continuous experience: stronger responses in DMN are observed when participants watch event changes in movies or listen to event changes in narratives (Baldassano et al., 2017; Swallow et al., 2011; Zacks et al., 2010). These various lines of evidence implicate AG in event processing, supporting the idea that this region may act as a semantic hub for event knowledge. If this is the case, it should represent semantic similarities between abstract event concepts (e.g., wedding-party), and it should code event similarities more strongly than object similarities. These predictions have not previously been tested directly.

In summary, vATL has emerged as a representational hub for various aspects of semantic knowledge, and is known to play an important role in coding similarity-based relationships between individual concepts. It is not clear whether this role extends to coding semantic relationships between more complex event concepts. In contrast, AG has been proposed to be a semantic hub that specialises for representing event-based knowledge, by integrating contextual information, interactions, and associations between objects. However, while numerous studies have investigated how this region responds to processing temporally-extended events (e.g., movies or narratives; Baldassano et al., 2017; Bonnici et al., 2016; Swallow et al., 2011; Zacks et al., 2010), it is less clear to what extent this region represents more abstract event concepts, or whether it represents these in preference to object concepts. More generally, the regions involved in representing semantic relations for objects and events have rarely been directly compared.

To address these questions, we used fMRI to scan participants when they were presented with event and object concepts (as written words and still images), then conducted RSA to test whether neural patterns reflected semantic similarity within either set of concepts. We particularly focused on representation similarity effects in vATL and AG, since these have been proposed as core semantic hubs for objects and events respectively. We analysed left and right vATLs and AGs. Many studies have assumed semantic representations are left-lateralised and have not tested effects in right-hemisphere regions. Here we included both hemispheres, to determine whether effects are specific to the left hemisphere. In addition, univariate analysis was conducted to test general activation differences to event and object concepts. Finally, psychophysiological interaction (PPI) analyses were performed to explore whether, when processing event and object concepts, semantic hubs have different connective patterns with other areas.

## Method

### Participants

We recruited 43 healthy participants (31 females, 12 males; mean age = 23.07 years, s.d. = 3.23 years, range = 19–32). All participants were right-handed native English speakers, and no-one reported history of dyslexia or other neurological disorders. The study was approved by University of Edinburgh School of Philosophy, Psychology & Language Sciences Research Ethics Committee.

### Materials

We presented participants with 60 different concepts, each of which was represented by four different pictures (240 pictures in total; see Figure 1A for examples). 30 of these were event concepts, while the other 30 were object concepts. The list of all concepts can be found in Supplementary Materials. Object concepts referred to individual entities, and we sampled from a variety of categories: animals (e.g., a dog), food (an apple), manipulable tools (a hammer), vehicles (a car), buildings (a castle), body parts (an arm) and human entities (a woman). Event concepts referred to situations in which multiple people or entities interact, including a range of social (e.g., a party), cultural (an opera), professional (a diagnosis) and everyday events (a picnic). In the experiment, each concept was presented 4 times, with the concept name shown each time with a different picture. We used images to elicit richer representations of the underlying concepts. In addition, by showing broader contexts and interactions, event pictures encouraged participants to process the situational aspects of these concepts. In contrast, object pictures included no background or interactions, encouraging people to process each object as an individual entity. In RSA analyses, we used the average neural responses across all 4 presentations of each concept. This ensured that the neural pattern for each concept represented general knowledge of the concept, rather than idiosyncratic features of one particular image.

**Figure 1.**
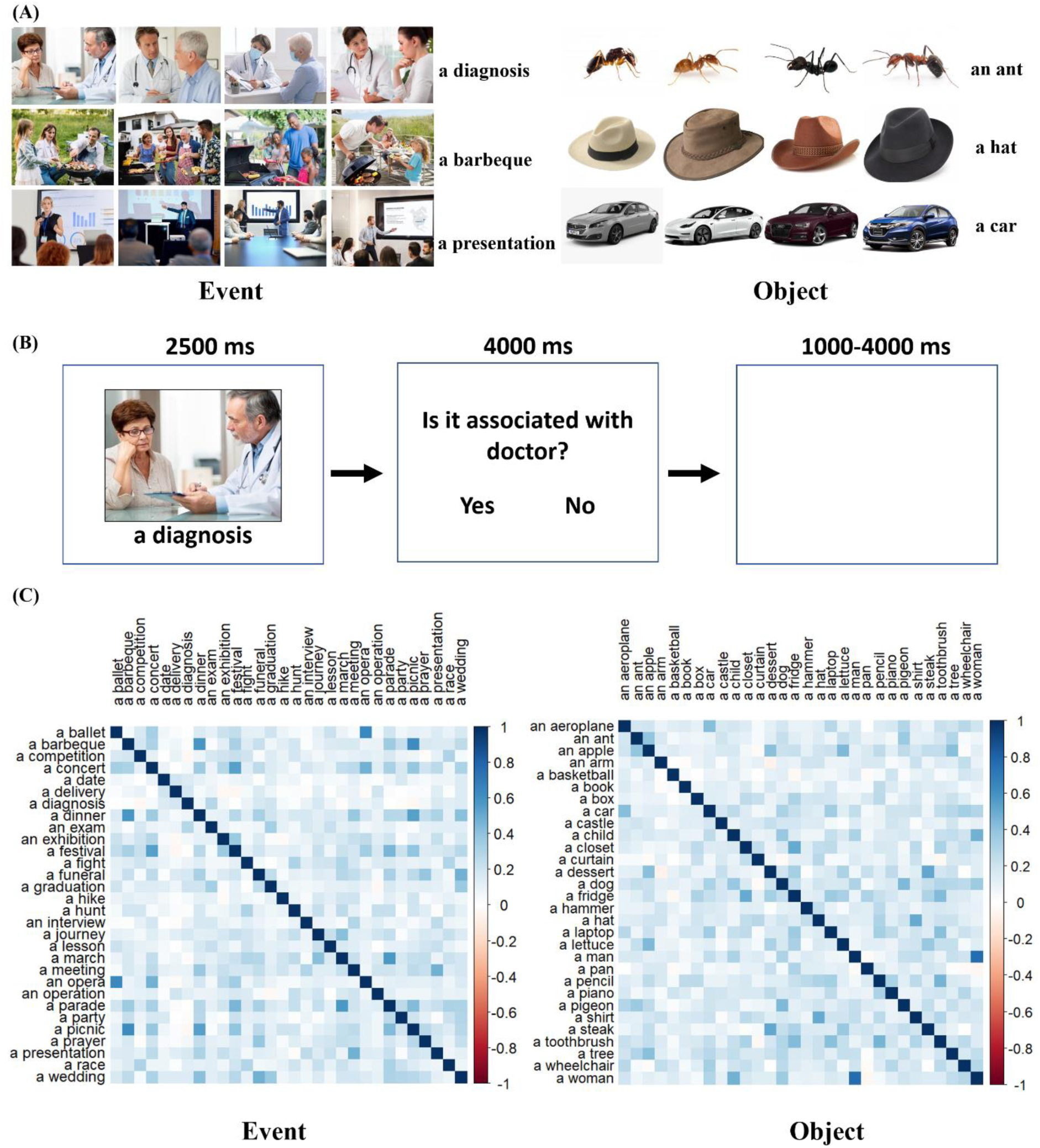
Experimental design.(A) Examples of object and event stimuli. (B) Experimental procedure, showing one trial followed by a catch question. (C) Semantic similarities for event concepts (left) and object concepts (right).

Object and event stimuli differed in several ways, reflecting intrinsic differences between object and event concepts. Compared with object pictures, event pictures were more visually complex because they showed scenes containing multiple people and objects. We also computed word frequency and concreteness for each concept name. Concreteness values were obtained from Brysbaert et al. (2014) and frequency values from Van Heuven et al. (2014). There was no significant difference between object and event concepts in frequency (*t*(58) = −0.04, p =0.97), but object concepts were more concrete than event concepts (*t*(58) = −9.95, *p* < 0.001). This was expected given that events are more complex and abstract than objects.

Given these differences, the main RSA analyses were conducted separately within each of the 2 sets of concepts. For completeness, we also present univariate activation contrasts of the two conditions but we note that effects in these contrasts could arise from differences at various levels of processing (i.e., lower-level visual perceptual processes as well as semantic processing).

For RSA, we constructed four 30*30 representational dissimilarity matrices (RDMs) that captured the similarity structures within events and within objects (see Figure 1C). For each set of concepts, we calculated a semantic RDM and a visual RDM. The semantic RDM was based on vector-based representations of word meaning, generated by training the word2vec neural network with the 100-billion word Google news corpus (Mikolov et al., 2013). We defined dissimilarity between two concepts as one minus the cosine between their word2vec vectors. Although a number of vector-based models of word meaning are available, we used word2vec because these vectors show the best fit to human semantic relatedness judgements (Pereira et al., 2016). The visual RDM controlled for the low-level visual characteristics of the images we presented. A visual representation of each image was calculated by entering images into the Hmax computational model of vision and extracting the output on the C1 layer of the model, which represents low-level visual attributes (Serre et al., 2007). Visual dissimilarity between images was defined as one minus the Pearson’s correlation between their C1 outputs (for a similar approach, see Naspi et al., 2021). To determine the visual dissimilarity between concepts, we averaged the pairwise dissimilarities between the images representing each concept.

### Experimental procedure

Participants viewed the concepts in a single scanning run of approximately 24 minutes, after completing two runs of unrelated tasks described later. The timeline for a single trial is shown in Figure 1B. Each trial consisted of a picture presented in the middle of screen for 2.5s with the concept name shown below. Participants were asked to think about the concept demonstrated by the picture and described by the word. To ensure that participants paid attention to the concepts, on 25% of trials the concept was followed by a catch question, which asked if the concept is related to another word. For example, for the concept ‘a diagnosis’, the catch question was ‘Is it associated with doctor?’. Each concept was followed by a catch question on one of its four presentations. The correct answers for half of these catch questions were ‘Yes’, for the other half they were ‘No’. All trials were presented with a mean interstimulus interval of 2.5 s, jittered between 1s and 4s. Trials were presented in 4 blocks, each containing one instance of each concept. The order of stimuli within each block was randomised separately for each participant.

### Image acquisition and processing

Images were acquired on a 3T Siemens Prisma scanner with a 32-channel head coil. For the functional images, the multi-echo EPI sequence included 46 slices covering the whole brain with echo time (TE) at 13 msec, 31 msec and 50 msec, repetition time (TR) = 1.7 sec, flip angle = 73, 80 * 80 matrix, reconstructed in-plane resolution = 3 mm * 3 mm, slice thickness = 3.0 mm (no slice gap) and multiband factor = 2. A single run of 858 volumes was acquired. A high-resolution T1-weighted structural image was also acquired for each participant using an MP-RAGE sequence with 1 mm isotropic voxels, TR = 2.5 sec, TE = 4.4 msec. To minimize the impact of head movements and signal drop out in the ventral temporal regions (Kundu et al., 2017), the study employed a whole-brain multi-echo acquisition protocol, in which data were simultaneously acquired at 3 TEs. Data from the three-echo series were weighted and combined, and the resulting time-series were denoised using independent components analysis (ICA).

Images were pre-processed and analysed using SPM12 and the TE-Dependent Analysis Toolbox (Tedana) (Kundu et al., 2013; Kundu, Inati, Evans, Luh, & Bandettini, 2012). Estimates of head motion were obtained using the first BOLD echo series. Slice-timing correction was carried out and images were then realigned using the previously obtained motion estimates. Tedana was used to combine the three-echo series into a single-time series and to divide the data into components classified as either BOLD-signal or noise-related based on their patterns of signal decay over increasing TEs (Kundu et al., 2017). Components classified as noise were discarded. After that, images were unwrapped with a B0 field-map to correct for irregularities in the scanner’s magnetic field. Finally, functional images were spatially normalised to MNI space using SPM’s DARTEL tool (Ashburner, 2007), and were smoothed with a kernel of 8 mm FWHM for univariate and PPI analysis and 4 mm FWHM for RSA analysis. Data in our study were treated with a high-pass filter with a cut-off of 180s. Covariates consisted of six motion parameters and their first-order derivatives.

For univariate and PPI analysis, a general linear model (GLM) was used that included 3 regressors for event concepts, object concepts, and catch trials. For RSA, to obtain better estimates of activation patterns of each concept, we used the least squares separate (LSS) approach (Mumford et al., 2012). We ran a separate GLM for each concept, where the 4 trials of that concept were modelled as the regressor of interest and all other trials were combined into a single nuisance regressor (with a further regressor modelling catch questions). This process yielded one activation map for each concept, which were used to compute neural RDMs.

### Regions of Interest

We defined 4 regions of interest (ROIs): left ventral anterior temporal lobe (left vATL), left angular gyrus (left AG), right ventral anterior temporal lobe (right vATL), right angular gyrus (right AG). Each ROI was defined as a 10mm radius sphere centred on specific MNI co-ordinates, which were selected in a two-stage process.

In the first stage, we constructed anatomical masks covering the vATLs and AGs. Masks of vATLs were made in a similar way to Hoffman and Lambon Ralph (2018). We first created masks of the temporal regions: inferior temporal gyrus, fusiform gyrus, superior temporal gyrus, and middle temporal gyrus. These were created by including all voxels with a greater than 50% probability of being located within these areas in the LONI Probabilistic Brain Atlas (LPBA40) (Shattuck et al., 2008). Then we divided each of these masks into 6 sections of roughly equal length along an anterior-to-posterior axis. These sections were numbered 0-5, with section 0 representing the most anterior section. The divisions were made approximately perpendicular to the long axis of the temporal lobe. Finally, we created separate left vATL and right vATL masks by combining sections 1 and 2 of temporal regions’ masks in the left hemisphere and right hemisphere, separately. For masks of AGs, we included all voxels with a greater than 30% probability of being located within this particular brain region as defined by the LPBA40 atlas (Shattuck et al., 2008).

Within these large anatomical masks, we then sought the voxels that were most responsive to semantic processing, using the activation peaks from an independent semantic > non-semantic contrast in the same participants. In the scanning runs prior to the object/events task, participants completed a series of tasks which required them to match words based on similarities in color, size, general meaning and letters (for further details, see Wu & Hoffman, 2023). The judgements of color, size and general meaning all required access to semantic knowledge, while the letter similarity task did not. Based on these tasks, we made a semantic > non-semantic contrast at the group level and identified the peak co-ordinates within each anatomical mask. In the vATLs, the maximal response was in the left and right anterior fusiform region. The maximal AG response was in the ventral part of the AG mask, in the region of the temporoparietal junction. Each ROI was defined as a 10mm radius sphere centred on the peak semantic > non-semantic co-ordinates within each anatomical mask (see Figure 3). The centre coordinates were: left vATL [-36, −18, −30]; left AG [-51, −54, 15]; right vATL [33, −9, −39]; right AG [66, −45, 15]. These 4 ROIs were used in univariate, RSA, and PPI analyses.

### Behavioural analysis

For the behavioral data, we built one linear mixed effect (LME) model to predict accuracy for responses to catch questions of event and object concepts, and another one to predict reaction times. The analyses were conducted with R-4.0.3, and 3 packages: ‘lme4’, ‘effects’ and ‘afex’. In each LME model, concept type (event/object) was set as a fixed effect, and participant was set as the random effect with intercepts and random slopes for concept type.

### Univariate analysis

To compare activation for event concept and object concept conditions, both whole-brain analysis and ROI analyses were conducted with SPM12. The whole-brain analysis was corrected for multiple comparisons (*p* < 0.05) at the cluster level using SPM’s random field theory, with a cluster-forming threshold of *p* < 0.005. In ROI analyses, we extracted mean beta values in left vATL, left AG, right vATL, right AG in each condition, which represent activation relative to the implicit baseline (rest). Then a three-way repeated ANOVA analysis was done using R-4.2.2, to examine the effects of concept type (event/object), ROI (AG/vATL), hemisphere (left/right) and their interactions.

### Representational similarity analysis (RSA)

We used RSA to examine which brain areas are sensitive to similarity in event and object concepts’ semantic representations. CoSMoMVPA (Oosterhof et al., 2016) was used for these analyses.

To investigate effects across the brain, we used a searchlight analysis with a spherical searchlight with radius of 4 voxels. We extracted activation patterns for the 60 concepts, and computed pairwise dis-similarities (1 – Pearson correlation) between activation patterns for the event concepts and separately for the object concepts. Then the partial Spearman correlation between neural RDMs and semantic RDMs, controlling for effects of the visual RDMs, was computed. This process was repeated for all searchlights, resulting in two correlation maps, one for objects and one for events. These showed the degree to which neural similarities between concepts are predicted by their semantic similarity. We also computed a difference map by subtracting the 2 correlation maps, to check where neural patterns differed in their alignment with semantics for objects vs. events. Correlations were Fisher-z transformed for group-level analysis. We conducted ROI analysis in the same way but using neural patterns from the 4 spherical ROIs.

To test the significance of the semantic-neural correlations, we used a two-stage method to perform permutation tests (Stelzer et al., 2013). We first computed the correlation maps between semantic RDMs and neural RDMs 100 times for each participant, with random reshuffling of the labels in the semantic and visual RDMs each time. This process provided a distribution of expected correlations under the null hypothesis for each participant. Then we used a Monte Carlo approach to compute a null correlation distribution at the group level (over all participants). To do this, we randomly selected one null correlation map from each participant’s null distribution and averaged these to generate a group mean. This process was repeated 10,000 times to generate a distribution of the expected group correlation under the null hypothesis. In searchlight analyses, we entered the observed and null correlation maps into the Monte Carlo cluster statistics function of CoSMoMVPA to generate a statistical map corrected for multiple comparisons using threshold-free cluster enhancement (Smith & Nichols, 2009). These maps were thresholded at corrected *p* < 0.05. For ROI analyses, we used the position of the observed group correlation in the null distribution to determine the p-value (e.g., if the observed correlation was greater than 95% of correlations in the null distribution, the p-value would be 0.05).

### Psychophysiological interaction (PPI) analyses

PPI analysis is a functional connectivity method for investigating task-specific changes in the relationship between different brain regions’ activity (Friston et al., 1997). While functional connectivity analyses often consider the temporal correlations between different brain regions in all conditions (including the resting state), PPI concentrates on connectivity changes caused by experimental manipulations (Ashburner et al., 2014; Gitelman et al., 2003; O’Reilly et al., 2012). For this study, PPI analysis was conducted to examine which brain regions would show increased correlation with our ROIs when representing event concepts relative to object concepts, or vice versa. The PPI analysis for each seed region (left vATL, left AG, right vATL, right AG) was conducted using SPM12 and the gPPI toolbox (McLaren et al., 2012) with the following steps. First, the seed region was defined as described in the Region of Interest section above, and the BOLD signal time-series extracted using the first eigenvariate. Then, gPPI was used to create a GLM with the following regressors:

1. The signal in the seed region.
2. One regressor coding for each experimental effect of interest, including event concepts, object concepts and catch questions.
3. The interaction between the signal in the seed region and each experimental effect.
4. Head movement covariates as included in the main univariate analysis.

This model was used for testing differences between PPI regressors (i.e., changes in connectivity driven by concept type) in the whole brain. Results were corrected for multiple comparisons (*p* < 0.05) at the cluster using SPM’s random field theory, with a cluster-forming threshold of *p* < 0.005.

## Results

### Behavioural data

LME models were used to test whether participants responded differently to catch questions about event and object concepts. There were no significant differences in accuracies between concept types (event M = 97.44%, SD = 0.04, object M = 96.98%, SD = 0.04, *z* (42) = 21.79, *p* = 0.29) and overall accuracy was very high, suggesting participants maintained attention through the experiment. Participants responded slightly faster to event questions (event M = 1.26 s, SD = 0.27 s, object M = 1.30 s, SD = 0.26 s, *t* (1815) = −2.152, *p* < 0.03).

### Univariate fMRI analysis

We began by contrasting activation to events and objects. While these results showed which regions are differentially engaged by the conditions, it is important to note that there were substantial visual differences in the stimuli used in each condition. Thus, these results may reflect both semantic and visual differences between event and object trials. The whole-brain analysis contrasting event and object concepts is displayed in Figure 2. Event concepts elicited more activation than objects bilaterally in fusiform gyrus, middle occipital gyrus and lingual gyrus, as well as anterior and posterior parts of superior and middle temporal gyri, hippocampus and parahippocampal regions, parts of the ventromedial prefrontal cortex and posterior cingulate. Higher activation in visual and scene-processing areas (e.g., parahippocampal gyrus and posterior cingulate) may reflect differences in the images used in the two conditions. Event images were more visually complex, contained a higher number of objects and included contextual elements not present in the object images (see Figure 1 for examples). Stronger responses to events in ventromedial prefrontal cortex and temporal pole could be due to the relevance of social interactions to events (Binney & Ramsey, 2020). Comparatively, object concepts elicited higher activation bilaterally in supramarginal gyrus (SMG), superior parietal cortex and parts of the dorsolateral prefrontal cortices.

**Figure 2.**
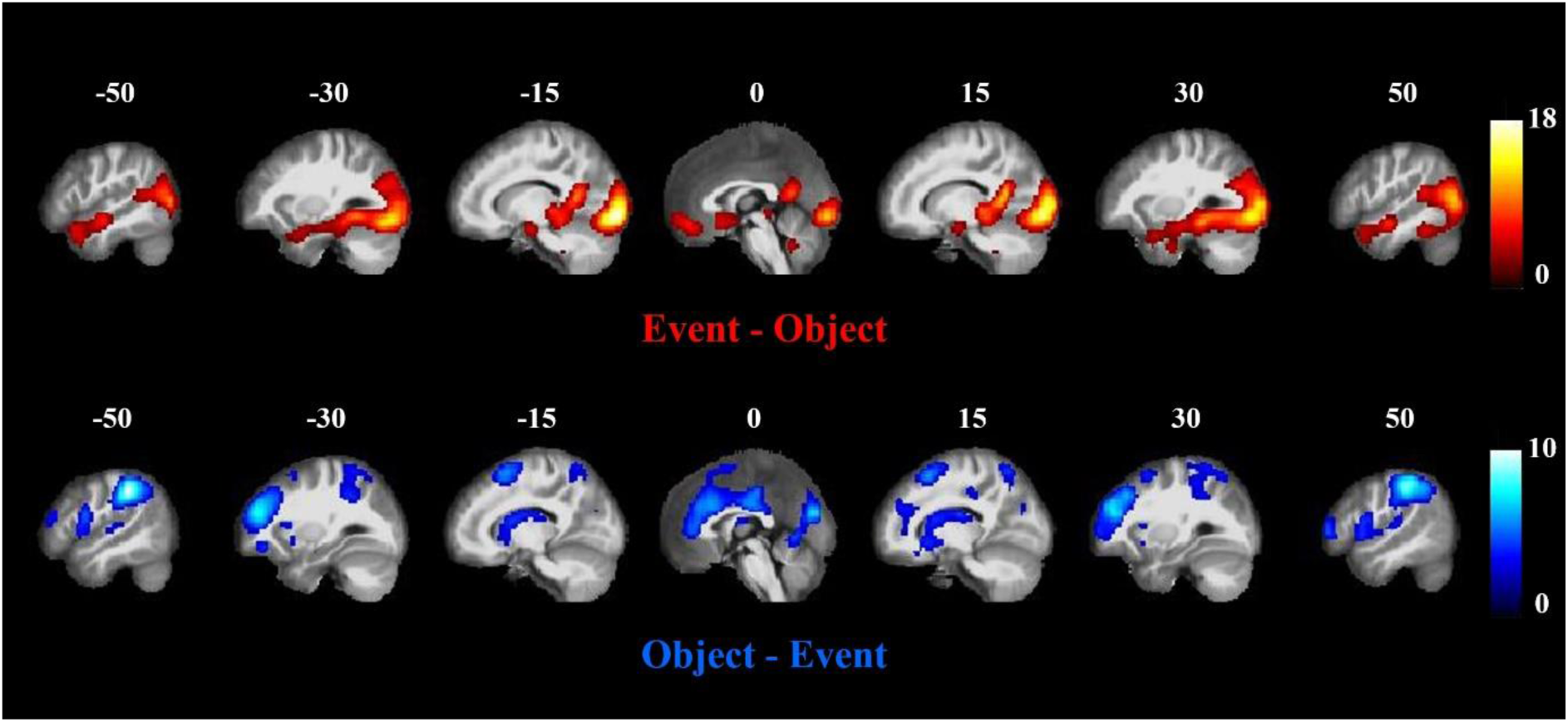
Univariate effects of event concepts versus object concepts, FWE corrected (p<0.05)

Figure 3 shows whether ROIs’ activations were affected by 3 factors: concept type (event/object), ROI (vATL/AG), hemisphere (left/right). A three-way repeated ANOVA was used to examine these effects. For both event and object concepts, ROIs in left hemisphere showed significantly higher activation (*F* (1, 42) = 15.88, p < 0.001). Overall, events elicited more activation than objects, and an interaction between concept type and ROI was also found (Concept effect: *F* (1, 42) = 4.436, *p* = 0.041; Concept x ROI: *F* (1, 42) = 5.483, *p* = 0.024). No other effects were significant. Post-hoc tests were performed comparing events vs. objects in each ROI. Left vATL was activated more strongly by events (*F* (1, 42) = 30.741, *p* <0.001), as was right vATL (*F* (1, 42) = 11.322, *p* = 0.002). There were no effects of concept type in left AG and right AG. According to dual-hub theory, vATL would be more engaged in processing objects, while AG is more engaged by event representation. The ROI analysis did not show this pattern. However, given the greater complexity of the event images, it is difficult to draw conclusions from these univariate analyses. For example, event images include multiple objects which could drive greater activation in object-specialised regions. To avoid this issue, we next conducted RSA within each concept type.

**Figure 3.**
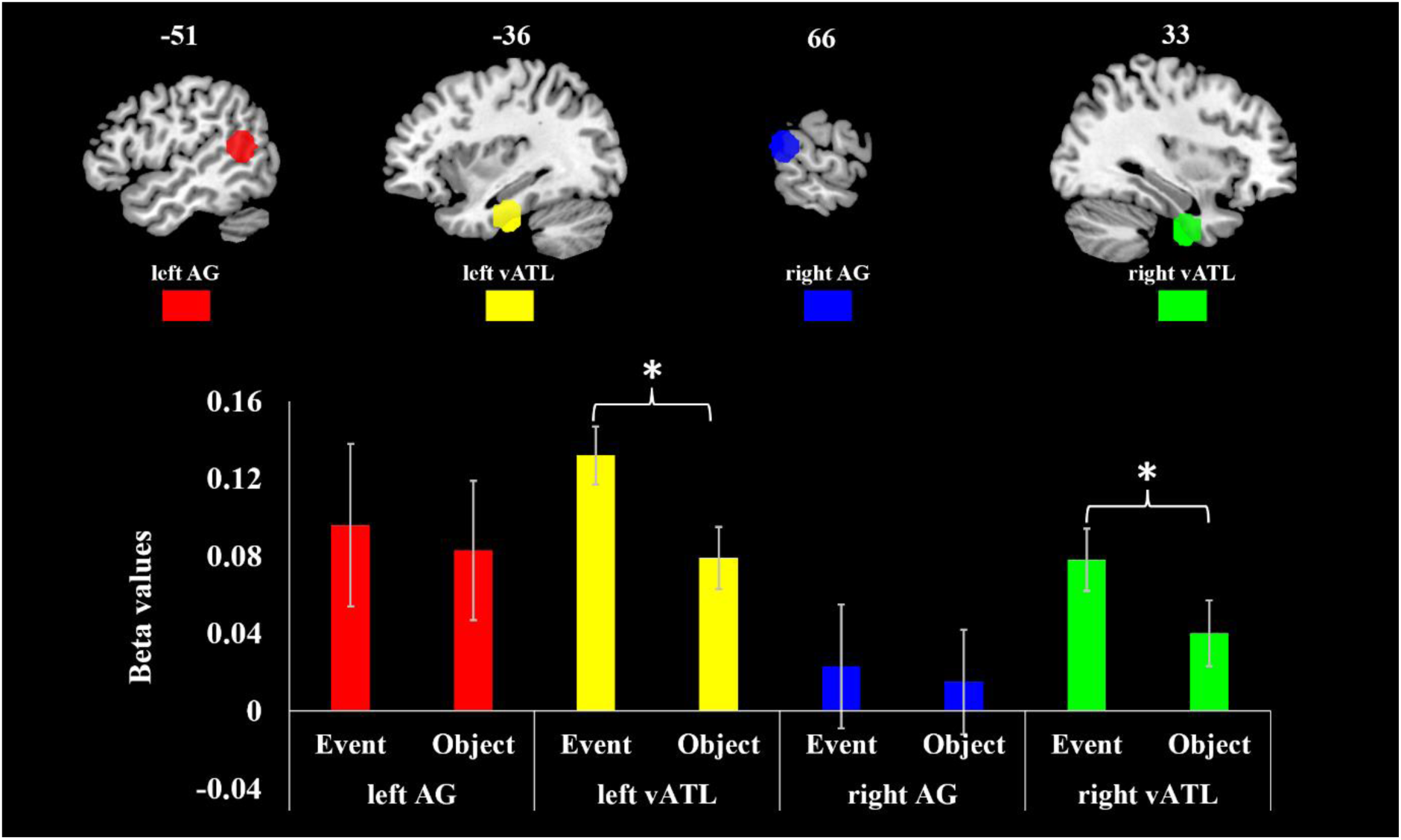
Activation to events and objects in ROIs. Bars show one standard error of the mean.

### Representational Similarity Analyses

The correlation maps, showing regions where neural RDMs were predicted by semantic RDMs, are displayed in Figure 4A. Generally, correlation effects were found in a similar set of bilateral regions for both events and objects. Specifically, the strongest effects were found in lateral occipital areas and parts of the ventral visual stream (ventral and medial temporal lobe), extending forward into vATL. We also observed effects spreading into TPC, especially for event concepts. The left inferior frontal area also showed correlations for both events and objects. Thus, neural activation patterns were correlated with semantic relationships not only in canonical semantic regions but also extensively in object and scene processing regions of the visual system. These effects indicate sensitivity to the semantic features of objects and events in these regions, since low-level visual similarity was controlled for in our analyses.

**Figure 4.**
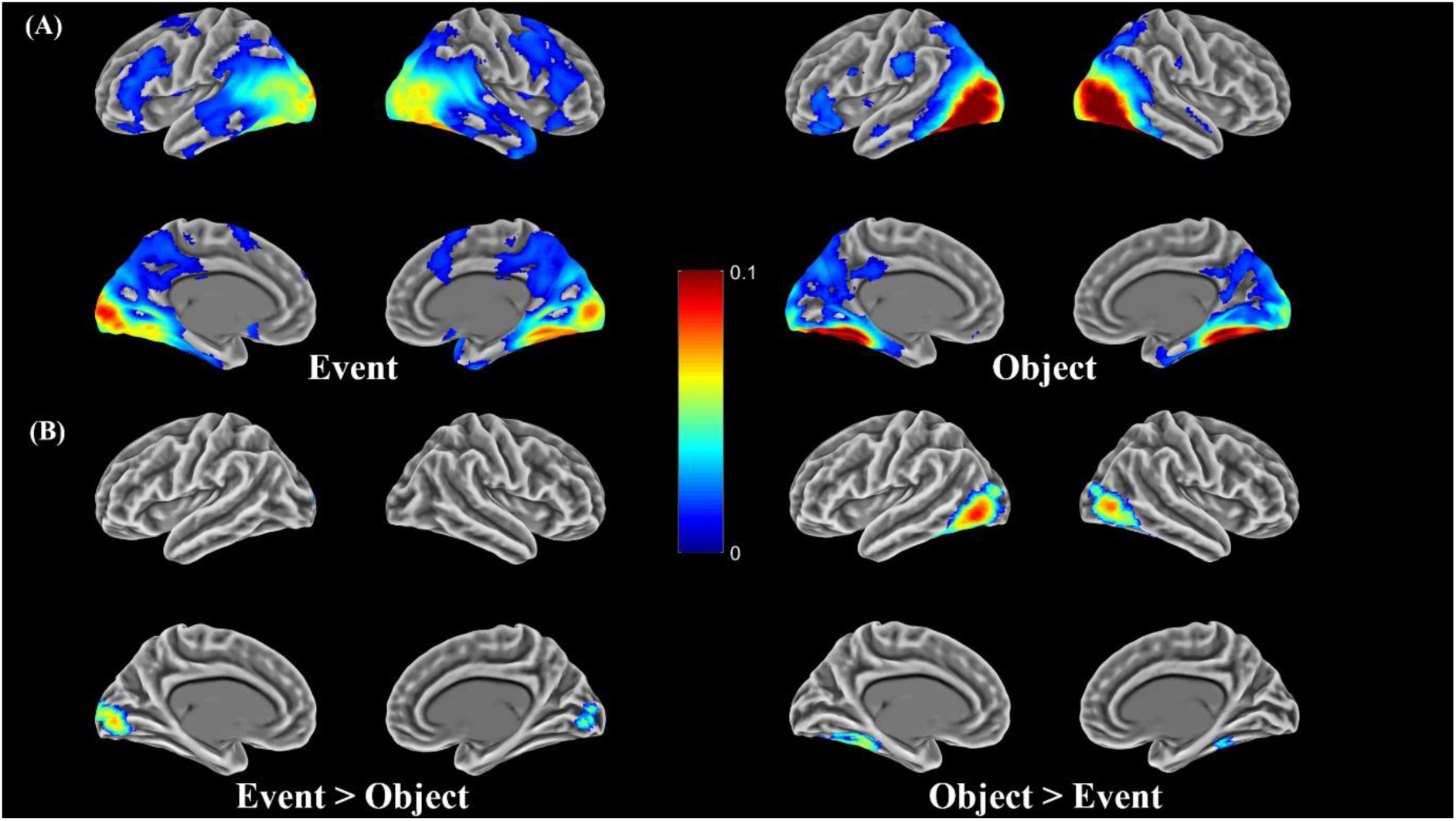
(A).Representational similarity maps for each concept type, showing regions where neural similarity is significantly correlated with semantic similarity (corrected p < 0.05); (B). The difference of representational similarities between event and object concepts (corrected p < 0.05). In (A) and (B), low-level visual features are controlled by covarying visual similarities measured with Hmax. Colour scale shows correlation strength.

Figure 4B presents regions that showed a significant difference in correlation strength between the event and object analyses. Bilateral primary visual cortex showed stronger correlations for events relative to objects. Conversely, stronger correlations for objects were found in lateral occipital regions, which is consistent with evidence for category-selective responses in this region in object recognition (for review, see Bi et al., 2016; Carota et al., 2017; Chen et al., 2017; Wu et al., 2020; Wurm & Caramazza, 2022). No differences were found in our target regions of vATL and AG, so we turned to more sensitive ROI analyses to investigate effects in these regions.

The correlations between neural and semantic RDMs in the four ROIs are displayed in Figure 5 Permutation testing indicated that left vATL, right vATL and left AG showed significant correlation between neural RDMs and semantic RDMs for both event and object concepts (all *p* < 0.0056). Right AG only showed a significant correlation for event concepts (*p* < 0.001).

**Figure 5.**
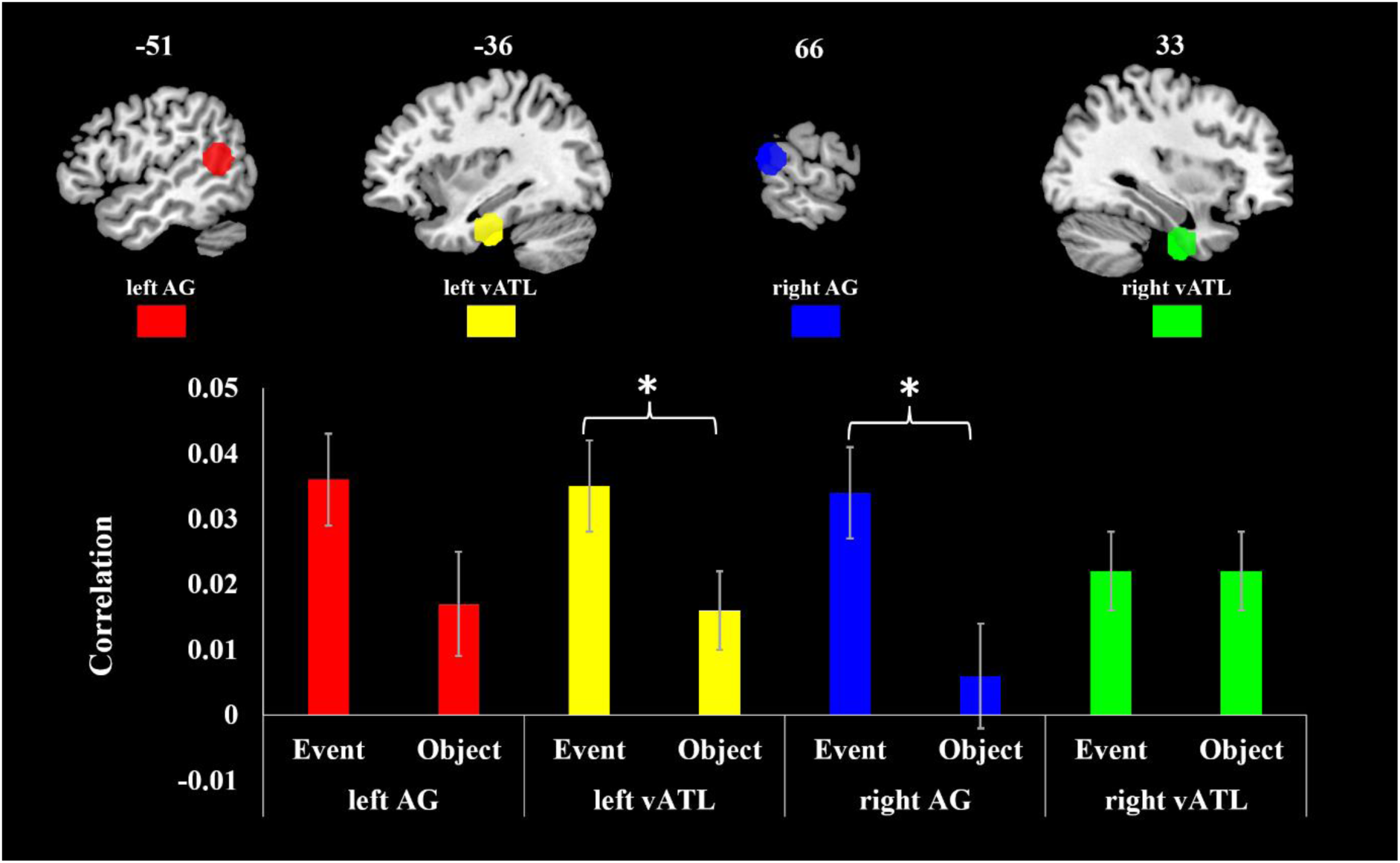
Representational similarity effects in ROIs

A three-way repeated ANOVA was conducted to examine whether correlations were affected by 3 factors: ROI (vATL/AG), hemisphere (left/right) and concept type (event/object). Overall, event concepts’ RDM showed higher correlations with neural RDMs than objects’ (*F* (1, 42) = 9.467, *p* = 0.004). No other main effects or interactions were significant at *p* < 0.05, though there was a suggestion of a weak three-way interaction (*F* (1, 42) = 3.27, *p* = 0.078). In post-hoc pairwise comparisons of events and objects in each ROI, left vATL and right AG had significantly higher correlations for event concepts (left vATL *F* (42) = 5.106, *p* = 0.03; right AG *F* (42) = 10.951, *p* = 0.002). Left AG also showed a stronger correlation for event concepts, but this difference was not statistically significant (*F* (42) = 3.362, *p* = 0.074). A two-way ANOVA (concept type*hemisphere) conducted on the AG data reported a main effect of concept type (*F* (1, 42) = 9.379, *p* = 0.004), but no interaction between concept type and hemisphere (*F* (1, 42) = 0.509, *p* = 0.479). This result suggests left AG and right AG had similar effects of concept type.

In a post-hoc two-way ANOVAs in data split by hemisphere (concept type*ROI), both left and right hemispheres showed significantly higher correlations for event concepts (left hemisphere *F* (1, 42) = 7.112, *p* = 0.011; right hemisphere *F* (1, 42) = 4.875, *p* = 0.033), and only right hemisphere showed interaction between ROI and concept type (*F* (1, 42) = 6.962, *p* = 0.012). This result suggests left vATL and left AG had similar effects of concept type, whereas right AG showed a stronger representational similarity for events than for objects compared to right vATL.

To summarise, stronger correlations for events than objects were found in bilateral AG and in left vATL. The results in AGs are consistent with the dual-hub hypothesis, which proposes that AG is specialised for representing semantic properties of events. However, effects in the vATLs contradict the idea that this region is particularly sensitive to object semantics. Our results instead indicate that right vATL is equally sensitive to events and objects’ semantics, while left vATL is more sensitive to events.

### Psychophysiological interaction (PPI) Analysis

To investigate how vATL and AG interact with other brain regions in representing concepts, PPI analyses were conducted using left vATL, left AG, right vATL and right AG as seed regions. Analyses tested for change in connectivity as a function of concept type (event vs. object). When participants processed event concepts, left vATL had stronger connectivity with right posterior MTG (Figure 6A). Right vATL showed a similar pattern but the effect did not survive cluster correction (see Supplementary Figure 1). Right AG showed stronger connectivity with bilateral fusiform gyrus and middle occipital gyrus (Figure 6B). Left AG showed no effects at cluster-corrected significance, though a more lenient uncorrected threshold showed increased connectivity with left fusiform gyrus, left ITG and right IFG for event concepts. Supplementary Figure 1 shows uncorrected events > objects effects for all four seed regions. No effects for objects > events were found at a cluster-corrected threshold and very few significant areas were found at an uncorrected threshold (shown in Supplementary Figure 2).

**Figure 6.**
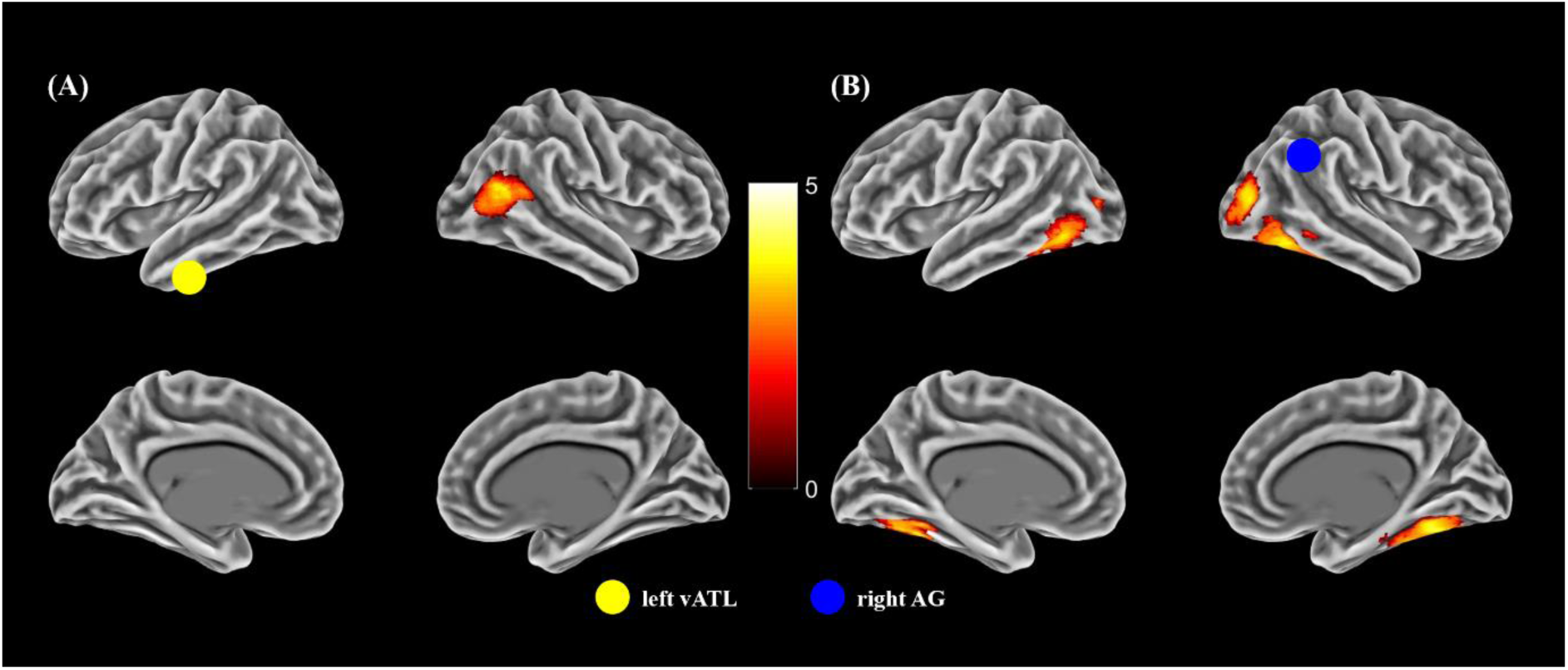
(A). For events > objects, regions showing increased connectivity with left vATL; (B). For events > objects, regions showing increased connectivity with right AG. Surface render (cluster corrected p<0.05). Seed regions are shown as coloured circles

## Discussion

Both event and object knowledge are critical semantic abilities, but their neural correlates are unclear. Some researchers have suggested that vATL is specialized for object semantics and AG for event semantics (Binder & Desai, 2011; Mirman et al., 2017). To test this hypothesis, we used RSA to investigate the neural basis of representing event and object concepts. Left and right AG were found to encode semantic similarity among event concepts more strongly than similarity among object concepts, though left AG also coded objects’ semantic similarity. Left and right vATLs both encoded semantic structure for object and event concepts, and left vATL showed stronger effects for events than objects. Univariate analyses also indicated more engagement of bilateral vATLs for event concepts. These findings support the idea that AG is more specialized for event semantics relative to object semantics. However, vATL specialization for object semantics is not supported by our results, suggesting that this region plays a more global role in semantic representation.

### Sensitivity to object and event semantics in the vATLs and AGs

Many previous studies have found that activity patterns in vATL code semantic similarities among object concepts (e.g., Bruffaerts et al., 2013; Chen et al., 2016; Clarke & Tyler, 2014; Devereux et al., 2018; Liuzzi et al., 2015; Naspi et al., 2021; Tyler et al., 2013). Our data indicate that the same region is also sensitive to semantic relationships between event concepts.

For vATLs, RSA indicated that their activity patterns reflect the semantic structure of events as well as objects (Figure 4), and left vATL showed a stronger correlation for events than objects. The simplest explanation for this is that vATL represents not only object, but also objects’ interactions and their context. The RSA finding is consistent with hub-and-spoke models of this region’s function (Lambon Ralph et al., 2017; Patterson et al., 2007; Rice, Hoffman, et al., 2015), which propose that vATL forms conceptual representations by integrating information from a range of neural sources. Our results suggest that, in addition to integrating the features of individual objects, this region may also form representations of more complex event-related concepts. However, an alternative explanation is that vATL *is* specialised for object representation and that the effects we see are a by-product of processing the objects involved in the depicted event stimuli. If semantically similar events involve semantically-similar objects, then vATL effects for events may reflect the coding for objects involved in those events. For example, *picnic* and *barbeque* are semantically similar events but they also contain semantically similar objects (food, plates, knives etc).

The univariate analysis showed more vATL activation for event trials (Figure 2). A possible explanation is that event concepts are more complex than object concepts, and therefore require greater semantic processing. According to hub-and-spoke theory, vATL integrates different modalities’ features into a concept, including not only visual features like color or shape, but also objects’ relevant actions or locations (Lambon Ralph et al., 2017; Peelen & Caramazza, 2012). Events contains multiple objects and people interacting in a specific environment. Thus, event concepts might lead the vATL to encode multiple concepts’ properties before settling on an overall representation of the event concept. The stronger vATL response for event concepts in univariate analysis might be caused by the heavier working load.

PPI analysis indicated that left vATL had stronger connectivity with right pMTG when processing event concepts (Figure 6A). Right pMTG has been implicated in coding causal relations between objects (Leshinskaya & Thompson-Schill, 2020), and in representing action concepts present in videos, still images and in language (Chen et al., 2020; Watson et al., 2013). The increased connectivity between vATL and pMTG may be a result of an enhanced contribution of relational and action-related information when understanding event concepts. This is in line with evidence that the vATL semantic hub alters its connectivity with more specialised spoke regions depending on the type of information that is relevant to the concepts being processed (Chiou & Ralph, 2019; Coutanche & Thompson-Schill, 2015).

For AG, RSA showed that activity patterns in both AG were correlated with semantic structure for events more strongly than for objects. Xu et al. (2018) also used RSA and found specialization of TPC for event-based relations among objects relative to category-based relations among the same objects. In contrast, the present study examined a single type of similarity (based on word2vec) and compared different types of concepts (events vs objects). Thus, the two studies provide converging complementary evidence of TPC (more specifically, AG) specialization for event semantics, consistent with region’s involvement in event representation more generally. AG plays an important role in representing autobiographical and episodic memories of events (Bonnici et al., 2018; Russell et al., 2019), in spatial-temporal feature integration (Ben-Zvi et al., 2015; Bonnici et al., 2016; Richter et al., 2016; Yazar et al., 2014; 2017), and in combinatorial semantics (Boylan et al., 2015). In addition, AG may be particularly sensitive to thematic relations because it processes contextual details of events (for review, see Binder & Desai, 2011; Mirman et al., 2017). AG is also part of the broader DMN, which integrates information to form context-specific representations of evolving situations (for review, see Ranganath & Ritchey, 2012; Yeshurun et al., 2021), and is sensitive to event boundaries within a continuous experience (Baldassano et al., 2017; Swallow et al., 2011; Zacks et al., 2010). These functions of AG together suggest that it encodes dynamic and complex combinations of concepts and experiences, where people, objects, and actions are bound together in time and space (for related proposals, see Humphreys & Lambon Ralph, 2015; Humphreys et al., 2021).

The univariate analysis did not show significant activation differences in AG between events and objects. This is not consistent with the idea that AG is specialised for event semantics. Bedny et al. (2014) used a similar univariate analysis and found stronger response in TPC (primarily posterior MTG) for event nouns relative to object nouns. A key difference between the two studies is that, in the present study, pictures were presented along with the nouns. Indeed, there were uncontrolled differences between event and object images, making these results (and differences from the results of Bedny et al.) hard to interpret.

Many previous studies implicating AG in event representation have presented temporally extended stimuli like narratives (e.g., Bonnici et al., 2016) or movies (e.g., Baldassano et al., 2017; Swallow et al., 2011; Zacks et al., 2010), or have required continuous generation of words (e.g., Bonnici et al., 2018; Yazar et al., 2014). In contrast, our study has shown that simple representations of static, abstract events are sufficient to engage AG for semantic processing. Furthermore, while previous language-based studies have focused on the role of left AG in representing thematic/event knowledge, here we found both left AG and right AG code event semantics (Figure 5). The bilateral effects might be due to our multimodal stimuli: while semantic activations are often left-lateralised for written word processing, more bilateral engagement is common for multimodal and non-verbal stimuli (Rice, Lambon Ralph, et al., 2015). Previous behavioural studies and lesion-symptom mapping studies indicated that left hemisphere injuries impaired verbal knowledge, while right hemisphere damage affected pictorial memory (Acres et al., 2009; Butler et al., 2009; Gainotti et al., 1994; Grossman & Wilson, 1987). Neuroimaging investigations further support this view, showing increased involvement of left temporal regions in processing verbal stimuli and right temporal cortex in understanding environmental sounds and images (Hocking & Price, 2009; Thierry et al., 2003; Thierry & Price, 2006).

In PPI analysis, right AG showed strong connectivity with bilateral ventral visual regions for event concepts (Figure 6B), which might be a consequence of this region extracting event-related information from the visual scenes we presented. An event image commonly incorporates a diverse set of agents and objects situated in a particular context. To represent an event as a cohesive concept, these individual items must be amalgamated, taking into account their identities, positions, orientations, and interactions. Increased connectivity between right AG and visual regions may reflect this process.

### Effects in other regions

In addition to the effects in vATL and AG, our RSA analysis also found that patterns throughout large portions of lateral and ventral occipitotemporal cortex (OTC) were correlated with semantic structure for both objects and events. Within these areas, correlations were stronger for object concepts than event concepts (Figure 4B). The correlation effects in OTC are consistent with selectivity for specific object categories in these regions (for review, see Bi et al., 2016). Many studies have reported that when people view pictures or object names, clusters of voxels in OTC are selectively responsive to certain categories of objects, such as faces, bodies, tools, or places (Chao et al., 1999; Costantini et al., 2011; Fairhall et al., 2014; Fairhall & Caramazza, 2013; Goyal et al., 2006; Ishai et al., 2000; Noppeney et al., 2006; O’Craven & Kanwisher, 2000). In particular, lateral OTC is known to be more strongly activated by small, manipulable objects (such as tools) and by body parts (Chao et al., 1999; Costantini et al., 2011; Noppeney et al., 2006). In ventral OTC, anterior medial regions (parahippocampal and medial fusiform) show preferences for inanimate items broadly related to navigation, including scenes, places, buildings, and large non-manipulable objects (Fairhall et al., 2014; Fairhall & Caramazza, 2013; Ishai et al., 2000; O’Craven & Kanwisher, 2000), while the posterior fusiform has a preference for animate items including faces and animals (Chao et al., 1999; Goyal et al., 2006; Ishai et al., 2000; O’Craven & Kanwisher, 2000). These category-selective responses explain why objects showed stronger semantic correlations with OTC patterns than events: objects from the same category were more semantically related, thus activated similar patches of cortex in OTC. Nevertheless, OTC patterns also showed correlations with event semantics. This could be because pictures of similar events tend to contain objects from similar categories, as discussed earlier.

Event concepts showed stronger correlations than object concepts in primary visual cortex. There are a few possible explanations for this effect. One intriguing possibility is that, when presented with static event images, participants were primed to mentally anticipate the movements of the objects or people depicted in those images. Primary visual cortex (V1) has been associated with motion-inducing illusion and predicting visual stimuli in many studies (Alink et al., 2010; Ekman et al., 2017; Gavornik & Bear, 2014; Kok et al., 2014; Muckli et al., 2005; Sterzer et al., 2006). V1 activation can be modulated by prediction of motion direction or onset (Alink et al., 2010) Muckli et al., 2005) and prior expectation of specific visual stimuli or visual sequences can evoke V1 responses similar to those evoked by viewing the actual stimuli or sequence (Ekman et al., 2017; Gavornik & Bear, 2014; Kok et al., 2014; Sterzer et al., 2006). For example, Ekman et al. (2017) found that after familiarizing participants with a spatial sequence, flashing only the starting point of the sequence triggered an activity wave in V1 that resembled the full stimulus sequence. Thus, the observed correlation effects in V1 might indicate the encoding of different predictions about potential motions in event images.

In conclusion, by testing the predictions of dual-hub theory with event and object concepts, our study found AG specialization for coding event semantics, but did not find vATL specialization for object semantics. Left vATL even coded similarity for events more strongly than objects. These findings provide new data on the divisions of labour that exist within the semantic system.

## Data Availability

Data and code supporting this study are available as follows. Neuroimaging data: https://doi.org/10.7488/ds/7521. Other data and analysis code: https://osf.io/mn8ft/. Group effect maps: https://neurovault.org/collections/NSWUEOPG/

## Credit authorship contribution statement

Yueyang Zhang: Conceptualization, Methodology, Investigation, Formal analysis, Writing – original draft. Wei Wu: Methodology, Investigation, Writing –review & editing. Daniel Mirman: Methodology, Writing –review & editing. Paul Hoffman: Funding acquisition, Conceptualization, Methodology, Formal analysis, Writing –review & editing.

## Acknowledgements

PH was supported by a BBSRC grant (BB/T004444/1). Imaging was carried out at the Edinburgh Imaging Facility (www.ed.ac.uk/edinburgh-imaging), University of Edinburgh, which is part of the SINAPSE collaboration (www.sinapse.ac.uk). We are grateful to the University of Minnesota Center for Magnetic Resonance Research for sharing their neuroimaging sequences. For the purpose of open access, the author has applied a Creative Commons Attribution (CC BY) licence to any Author Accepted Manuscript version arising from this submission.

## Supplementary Figures

**Supplementary Figure 1.**
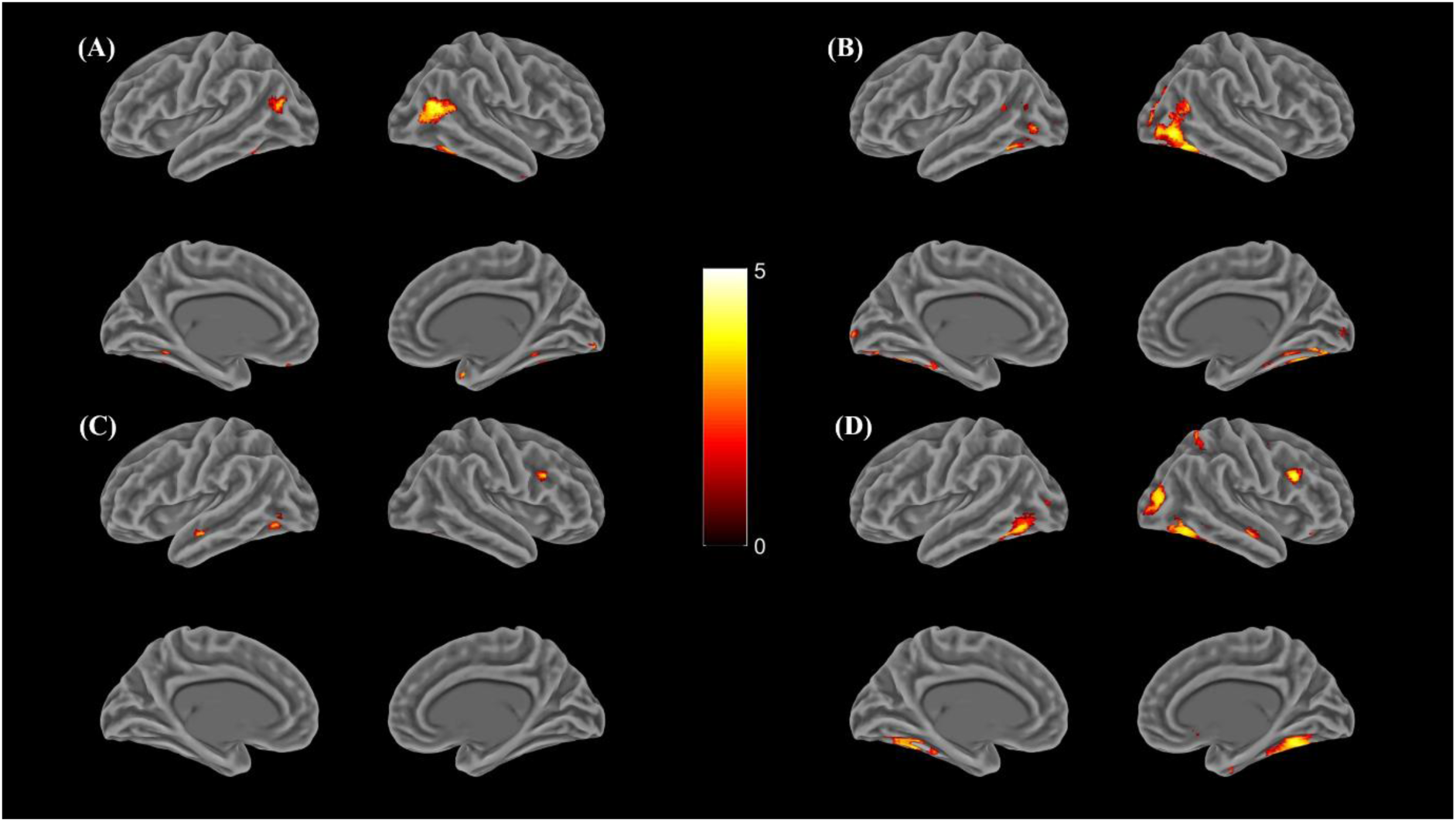
For events > objects, regions showing increased connectivity with (A). Left vATL; (B). Right vATL; (C). Left AG; (D). Right AG. Surface render (p<0.005, no cluster correction)

**Supplementary Figure 2.**
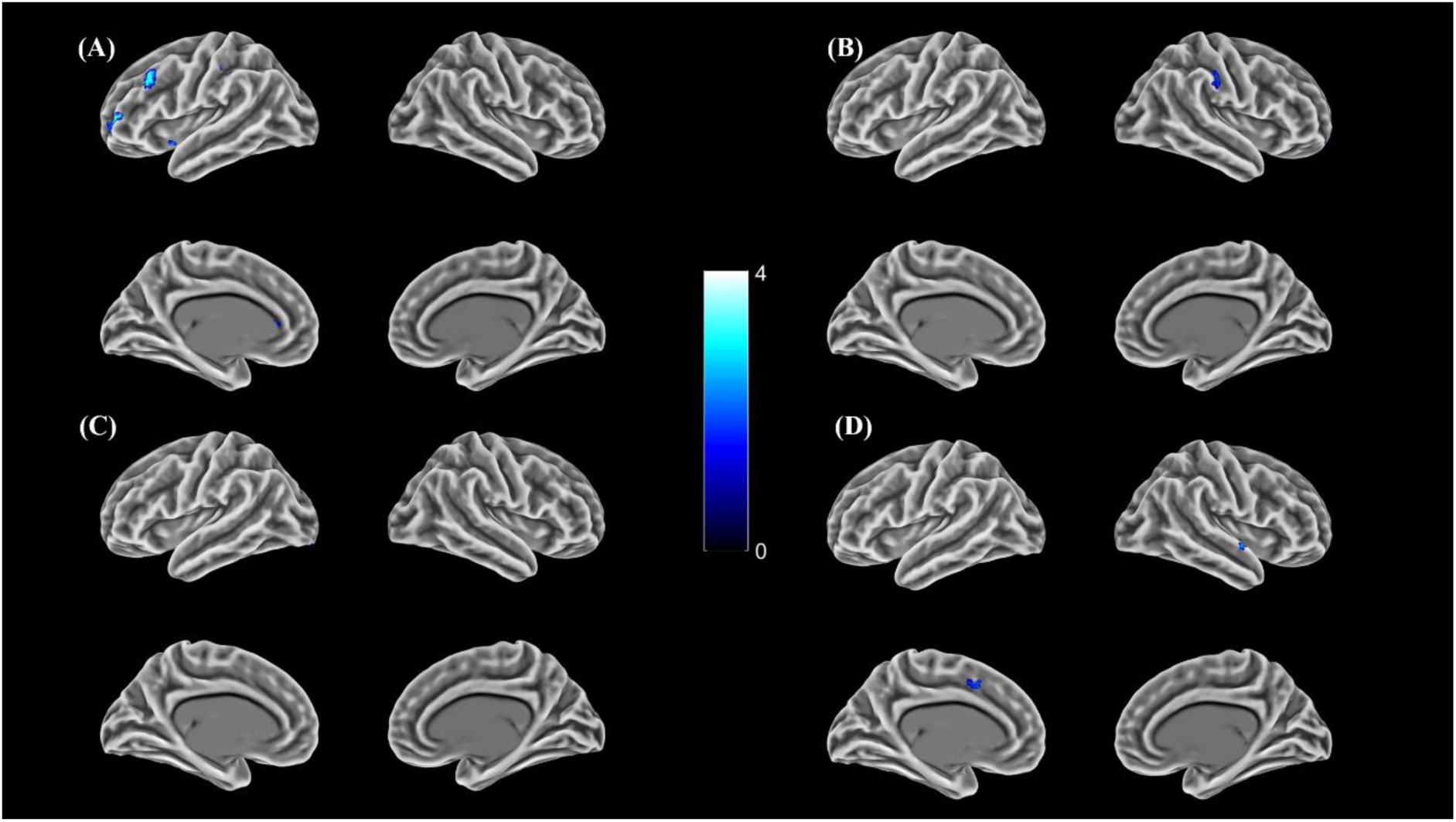
For object > events, regions showing increased connectivity with (A). Left vATL; (B). Right vATL; (C). Left AG; (D). Right AG. Surface render (p<0.005, no cluster correction)

